# Complete Protection of Nasal and Lung Airways Against SARS-CoV-2 Challenge by Antibody Plus Th1 Dominant N- and S-Specific T-Cell Responses to Subcutaneous Prime and Thermally-Stable Oral Boost Bivalent hAd5 Vaccination in an NHP Study

**DOI:** 10.1101/2020.12.08.416297

**Authors:** Elizabeth Gabitzsch, Jeffrey T. Safrit, Mohit Verma, Adrian Rice, Peter Sieling, Lise Zakin, Annie Shin, Brett Morimoto, Helty Adisetiyo, Raymond Wong, Ashish Bezawada, Kyle Dinkins, Joseph Balint, Victor Peykov, Hermes Garban, Philip Liu, Pete Bone, Andrew Bacon, Jeff Drew, Daniel C. Sanford, Patricia Spilman, Lennie Sender, Shahrooz Rabizadeh, Kayvan Niazi, Patrick Soon-Shiong

## Abstract

We have developed a dual-antigen COVID-19 vaccine incorporating genes for a modified SARS-CoV-2 spike (S-Fusion) protein and the viral nucleocapsid (N) protein with an Enhanced T-cell Stimulation Domain (N-ETSD) with the potential to increase MHC class I/II responses. The adenovirus serotype 5 platform used, hAd5 [E1-, E2b-, E3-], previously demonstrated to be effective in the presence of Ad immunity, can be delivered in an oral formulation that overcomes cold-chain limitations. The hAd5 S-Fusion + N-ETSD vaccine was evaluated in rhesus macaques showing that a subcutaneous prime followed by oral boosts elicited both humoral and Th1 dominant T-cell responses to both S and N that protected the upper and lower respiratory tracts from high titer (1 x 10^6^ TCID_50_) SARS-CoV-2 challenge. Notably, viral replication was inhibited within 24 hours of challenge in both lung and nasal passages, becoming undetectable within 7 days post-challenge.

**ONE SENTENCE SUMMARY:** hAd5 spike + nucleocapsid SC prime/oral boost vaccine elicits humoral and T-cell responses and protects rhesus macaques from SARS-CoV-2 challenge.

## INTRODUCTION

To address the ongoing COVID-19 pandemic (*1, 2*), particularly in the face of viral evolution and evidence of viral variant resistance to antibodies and convalescent plasma (*3-6*), we have developed a vaccine anticipated to protect individuals from SARS-CoV-2 that has the potential to not only elicit robust humoral responses but also activate T cells. The dual-antigen hAd5 S-Fusion + N-ETSD vaccine (Supplementary Fig. 1) expresses a viral spike (S) protein (S-Fusion) fused to a signal sequence that, as predicted based on reports for similar sequences (*7, 8*), in our *in vitro* studies enhances cell-surface expression of the spike receptor binding domain (S RBD) as compared to S wildtype (Supplementary Fig. 2), the antigen used in the majority of other vaccines being developed. Our vaccine also expresses the viral nucleocapsid (N) protein with an Enhanced T-cell Stimulation Domain (N-ETSD) that directs N to the endo/lysosomal subcellular compartment as demonstrated in Sieling *et al*. (in preprint) (*9*) which is predicted to enhance MHC class II responses (*10-12*).

**Fig. 1.**
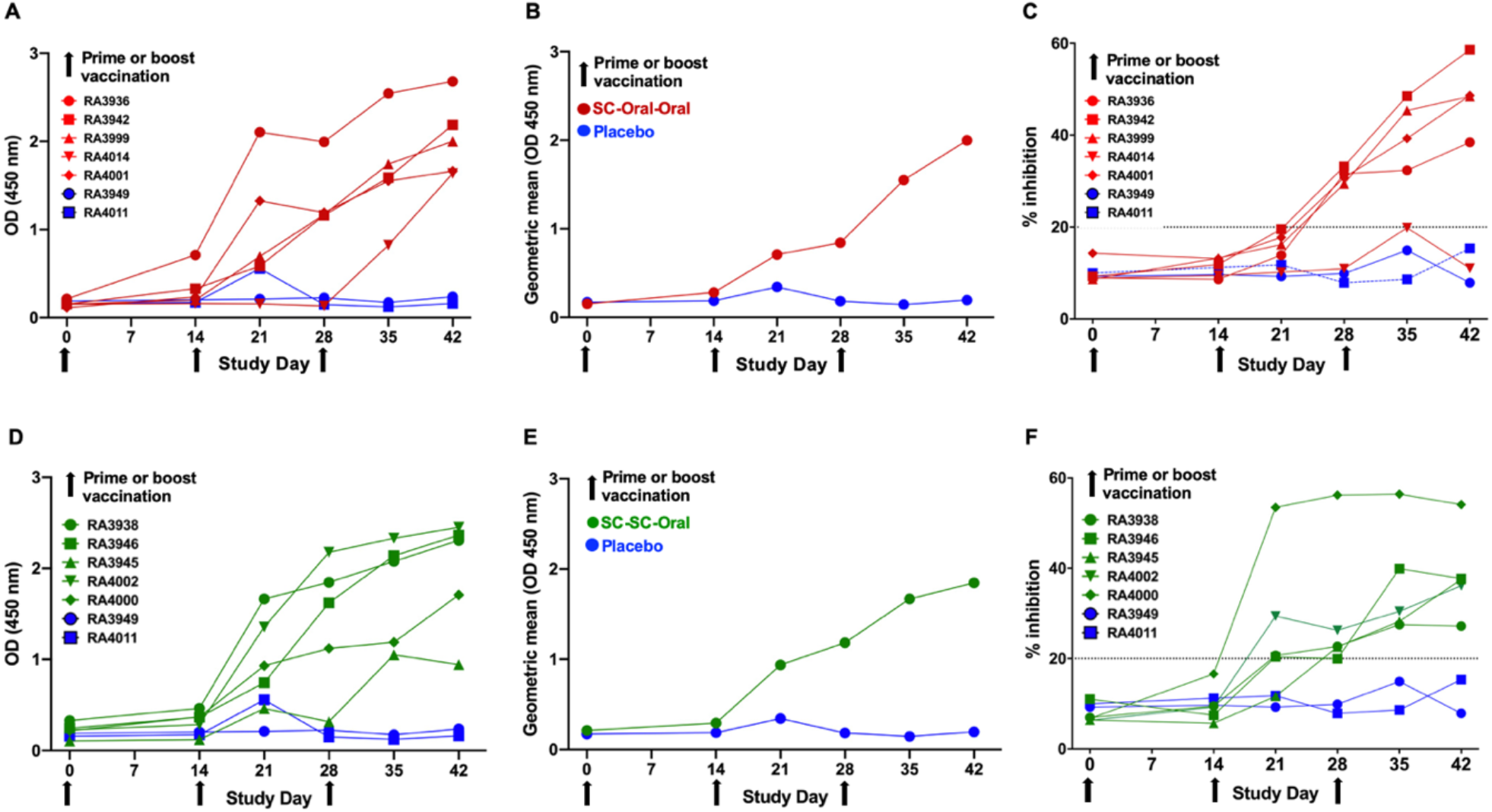
Humoral responses and neutralizing capability of sera from hAd5 S-Fusion + N-ETSD vaccinated NHP. Anti-spike IgG levels (ELISA; OD 450nm) are shown for (A) individual SC-Oral-Oral NHP along with the (B) geometric mean and (C) inhibition in the surrogate assay. These data are also shown for SC-SC-Oral NHP: (D) individual anti-S IgG, (E) geometric mean, and (F) inhibition in the surrogate assay. Inhibition above 20% (dashed line) with a sera dilution of 1:30 is correlated with neutralization of SARS-CoV-2 infection. NHP received vaccination on Days 0, 14 and 28 (black arrows).

**Fig. 2.**
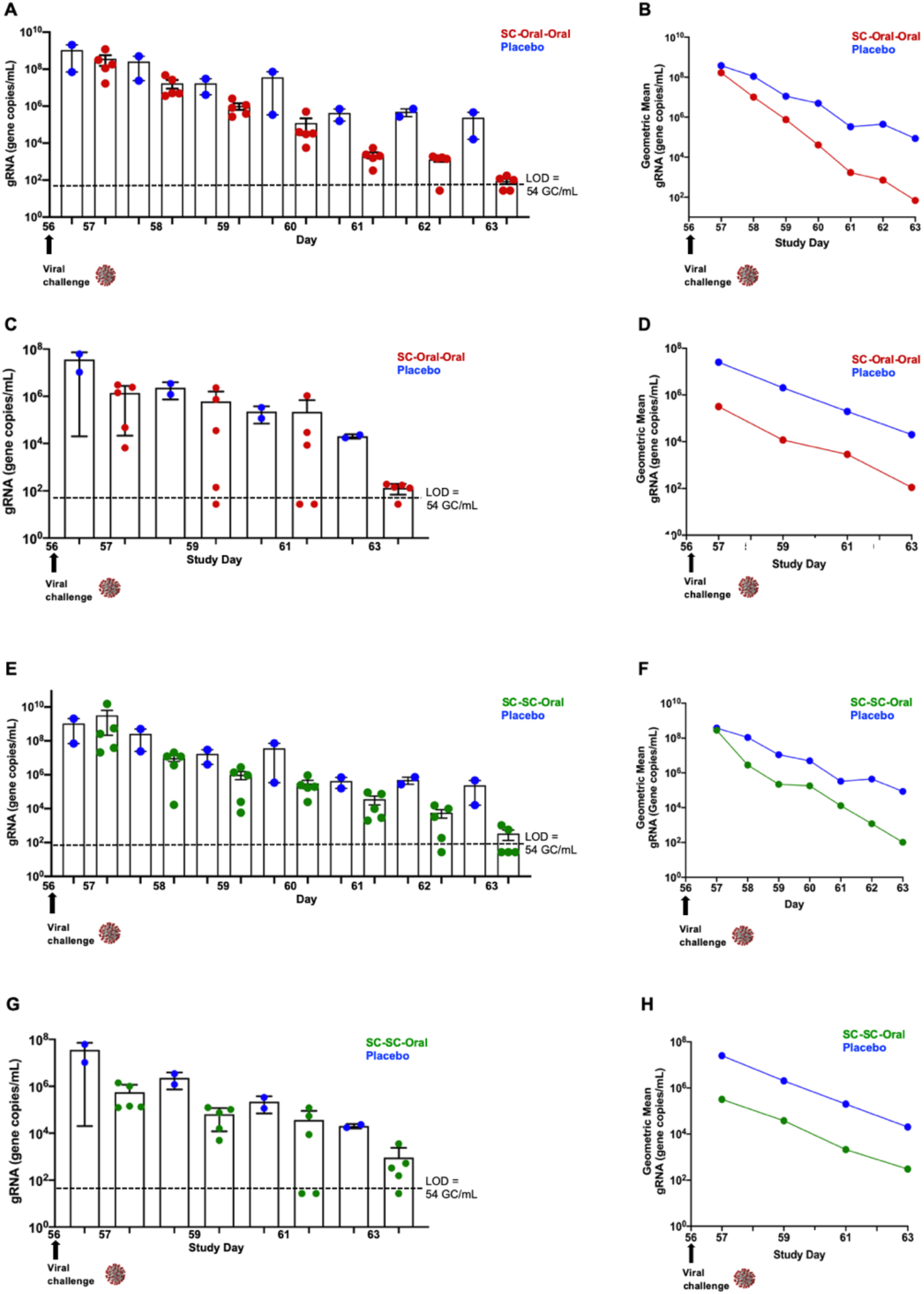
Viral load (gRNA) in nasal passages and lung of SC-Oral-Oral and SC-SC-Oral vaccinated NHP post-challenge. (A) Individual viral gRNA (RT qPCR) and (B) the geometric mean for nasal swab samples; and (C) gRNA and (D) the geometric mean for bronchoalveolar lavage (BAL) samples from SC-Oral-Oral NHP. (E) Individual gRNA and (F) the geometric mean for nasal swab samples; and (G) gRNA and (H) the geometric means for BAL samples from SC-SC-Oral NHP. SARS-CoV-2 challenge was on Day 56 (black arrows). The level of detection (LOD; dashed line) was 54 gene copies/mL (GC/mL) for gRNA and 119 GC/mL for sgRNA. For values below the LOD, half the LOD value (or 27 GC/mL for gRNA and 59 GC/mL for sgRNA) was used for graphing of individual values and calculation of the geometric mean.

The SARS-CoV-2 vaccine antigens are delivered by an recombinant human adenovirus serotype 5 (hAd5) [E1-, E2b-, E3-] vector platform (Supplementary Fig. 1A) we previously developed to rapidly generate vaccines against multiple agents, allowing production of high numbers of doses in a minimal time frame. The hAd5 platform has unique deletions in the early 1 (E1), early 2 (E2b) and early 3 (E3) regions (hAd5 [E1-, E2b-, E3-]), which distinguishes it from other adenoviral vaccine platform technologies under development (*13-15*), and allows it to be effective in the presence of pre-existing adenovirus immunity (*16-19*). We have utilized this platform to produce vaccines against viral antigens such as Influenza, HIV-1 and Lassa fever and have shown induction of both antibodies and cell mediated immunity (*18, 20-23*). In 2009, we showed that vaccination of mice with the hAd5 [E1-, E2b-, E3-] vector expressing H1N1 hemagglutinin and neuraminidase genes (*20*) elicited both cell-mediated immunity and humoral responses that protected the animals from lethal virus challenge.

The overwhelming majority of other SARS-CoV-2 vaccines in development target only the wildtype S antigen and are expected to elicit SARS-CoV-2 neutralizing antibody responses. In the development of our vaccine, we prioritized the activation T cells to enhance the breath and duration of protective immune responses (*24*); the addition of N in particular was predicted to afford a greater opportunity for T cell responses (*25-27*). T cells may provide immune protection at least as important as the generation of antibodies. In a study of SARS-CoV-2 convalescent patients, virus-specific T cells were seen in most patients, including asymptomatic individuals, even those with undetectable antibody responses (*27*).

In our preliminary studies of the hAd5 S-Fusion + N-ETSD vaccine in a murine model, we saw that the vaccine not only elicits T helper cell 1 (Th1)-dominant antibody responses to both S and N, it activates T-cells (Supplementary Fig. 3). We then looked the potential of the hAd5 S-Fusion + N-ETSD vaccine to re-capitulate T-cell activation prompted by natural SARS-CoV-2 infection (*9*). To do this, we transduced monocyte-derived dendritic cells (MoDCs) from SARS-CoV-2 convalescent individuals with the dual-antigen vaccine, incubated the S-Fusion and N-ETSD-expressing MoDCs with T cells from those same individuals, and saw that the vaccine antigens induce interferon-γ (IFN-γ) secretion by both CD4+ and CD8+ T cells (9). This demonstrates that T cells from SARS-CoV-2 convalescent individuals ‘recall’ the S-Fusion and N-ETSD antigens presented by transduced MoDCs as if they were re-exposed to the virus itself. This T-cell recall of vaccine antigens suggests that, conversely, hAd5 S-Fusion + N-ETSD vaccination will generate T cells that will recognize SARS-CoV-2 antigens upon viral infection and protect the vaccinated individual from disease.

**Fig. 3.**
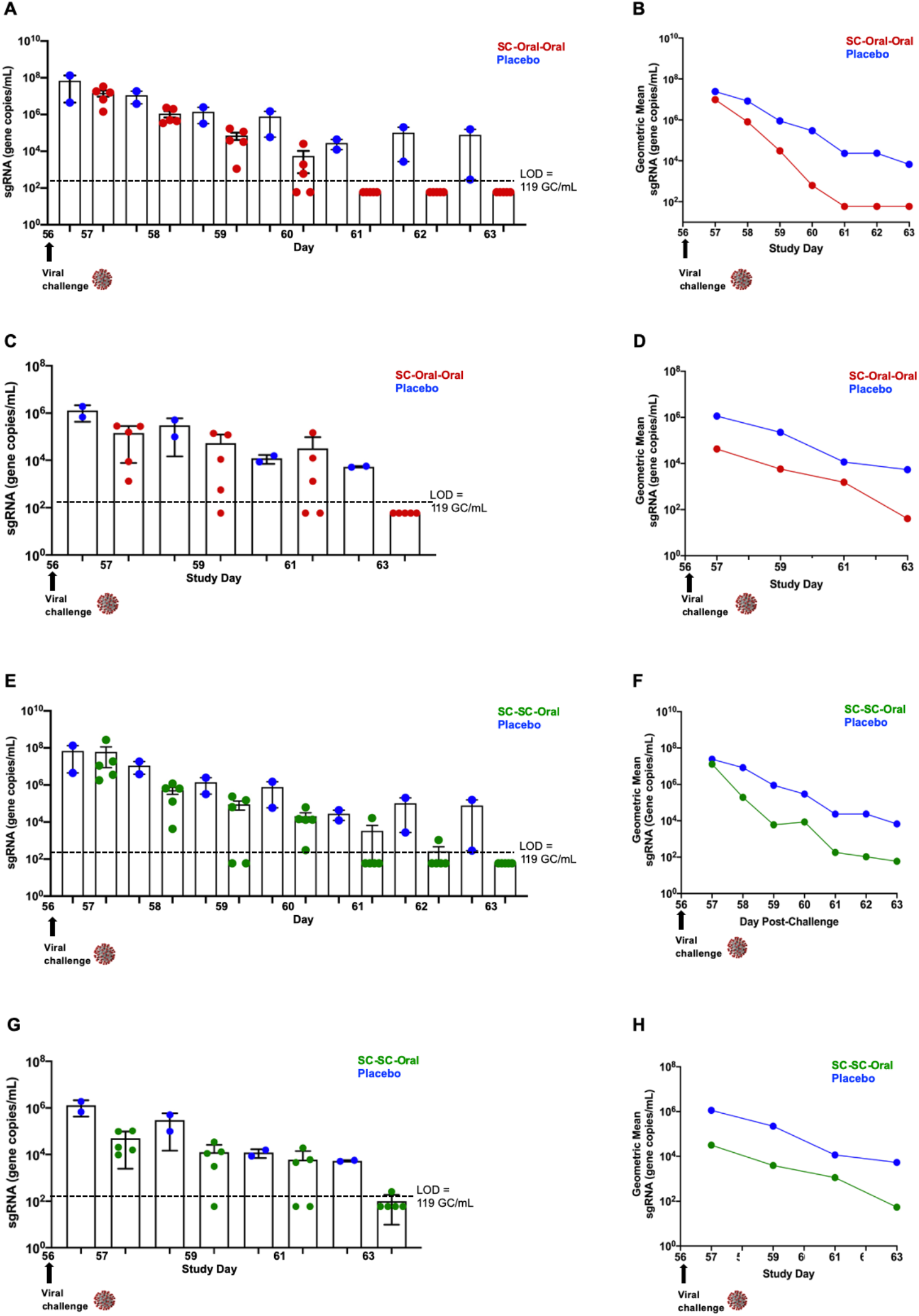
Viral replication (sgRNA) in nasal passages and lung in SC-Oral-Oral and SC-SC-Oral vaccinated NHP post-challenge. (A) Individual viral sgRNA (RT qPCR) and (B) the geometric mean for nasal swab samples; and (C) sgRNA and (D) the geometric mean for bronchoalveolar lavage (BAL) samples from SC-Oral-Oral NHP. (E) Individual sgRNA and (F) the geometric mean for nasal swab samples; and (G) sgRNA and (H) the geometric means for BAL samples from SC-SC-Oral NHP. SARS-CoV-2 challenge was on Day 56 (black arrows). The level of detection (LOD; dashed line) was 54 gene copies/mL (GC/mL) for gRNA and 119 GC/mL for sgRNA. For values below the LOD, half the LOD value (or 27 GC/mL for gRNA and 59 GC/mL for sgRNA) was used for graphing of individual values and calculation of the geometric mean.

The generation of T-cell responses may be a critical feature for a vaccine to be efficacious against the many variants whose emergence, at least in part, may be an escape response to antibodies generated by either first wave virus (*28*) or, as suggested in Wang *et al*. (in preprint) (*29*) by antibody-based vaccines. As further reported in Wang *et al*., neutralization by 14 of 17 of the most potent mRNA vaccine-elicited monoclonal antibodies (mAbs) was either decreased or abolished variants E484K, N501Y or the K417N:E484K:N501Y combination (*6, 30*). They also found these variants were selected for when recombinant vesicular stomatitis virus (rVSV)/SARS-CoV-2 S was cultured in the presence of these mAbs, which is highly suggestive that the presence of these antibodies act as an evolutionary force driving the appearance of new variants. T cells are not vulnerable to such forces and, if effectively established by vaccination, may provide protection (*31-33*) against existing viral strains and escape mutants.

We hypothesized that vaccination with the hAd5 S-Fusion + N-ETSD vaccine, which generates both neutralizing antibodies and memory T cells, would be protective in the rhesus macaque and that room temperature oral formulations would be effective as boosts.

In the next step in development of the hAd5 S-Fusion + N-ETSD vaccine, we formulated GMP (Good Manufacturing Practice)-grade liquid and oral forms of the vaccine for testing in non-human primates (NHP). A key objective of the NHP study design was to assess the efficacy of a subcutaneous (SC) prime followed by a thermally-stable (*34*) oral boost.

An oral boost provides several advantages in SARS-CoV-2 vaccination, including a greater potential for generating mucosal immunity particularly in the gastrointestinal tract (*35*) one of the major sites of infection (*36*). SARS-CoV-2 is a mucosal virus and is only rarely detected in blood (*37, 38*), therefore vaccines that specifically target mucosal immunity are of interest (*39*). Compelling additional advantages of a thermally-stable oral boost are that it would likely transform the global distribution of vaccines, especially in developing nations and potentially enable patients to self-administer the boost(s) at home. Because the hAd5 S-Fusion + N-ETSD construct induces both humoral and CMI responses to both antigens, it also has the potential to serve as a ‘universal’ heterologous booster vaccine to the multitude of SARS-CoV-2 vaccines under development.

Here, we report our findings from our study designed to determine the efficacy of the hAd5 S-Fusion + N-ETSD vaccine in rhesus macaques when delivered as either an SC prime with SC and oral boosts (SC-SC-Oral; n =5) or as an SC prime and two oral boosts (SC-Oral-Oral; n = 5) both using a regimen of prime on Day 0 and boosts on Days 14 and 28 to maximize T cell responses. Design details are presented in *Materials and Methods* and Supplementary Fig. S4. The goals of the study were to assess the immunogenicity of a dual-antigen hAd5 vaccine in both SC and oral formulations, and the potential of oral dose to serve as a boost following a single SC prime. Most importantly, we sought to assess cell-mediated T-cell response and protection of nasal passages and lung from SARS-CoV-2 infection after challenge as well as the rate of viral clearance.

### Clinical signs, hematology and clinical chemistry

No clinical signs were noted during the twice daily observations for clinical signs of toxicity due to vaccination and no animals died during the two weeks after one subcutaneous immunization of 1x 10^11^ vaccine particles (VP) or a week after an oral booster of 1×10^10^ IU of hAd5-S-Fusion+N-ETSD. In addition, no gross pathological effects or adverse events were observed and there were no notable changes in body weight (Supplementary Fig. 5). Lastly, hematology and clinical chemistry revealed no abnormalities as a result of vaccination (Supplementary Tables 1 and 2).

### An SC prime with oral boosts elicits generation of neutralizing, anti-spike antibodies

As shown in Figure 1, all SC-Oral-Oral vaccinated NHP produced anti-S IgG that increased after both the Day 14 and Day 28 oral boosts (Fig. 1A and B). Sera from 4 of 5 SC-Oral-Oral NHP, taken at baseline and every week starting at Day 14 and up to Day 42, demonstrated inhibition in the neutralization assay (Fig. 1C) that assesses the inhibition of binding of S RBD to recombinant angiotensin-converting enzyme 2 (ACE2) and is reported to correlate with the ability of sera to neutralize the SARS-CoV-2 virus (*40*).

Anti-S IgG production was similar for SC-SC-Oral (where the first boost was SC) hAd5 S-Fusion + N-ETSD vaccinated NHP (Fig. 1D and E) and sera from all five NHP in this group demonstrated inhibition in the surrogate assay for viral neutralization (Fig. 1F).

### SC prime, oral boost vaccination reduces viral load in nasal passages and lung after SARS-CoV-2 challenge

RT-qPCR analysis of genomic RNA (gRNA) was performed on nasal swab and bronchoalveolar lavage (BAL) samples to determine the amount of virus present. SC-Oral-Oral vaccination of NHP reduced SARS-CoV-2 gRNA in the nasal swab samples as compared to placebo control NHP from Day 57, the first day after challenge (Fig. 2A and B; Supplementary Table 3). Viral gRNA in this group continued to diminish to levels that were very low or below the level of detection (LOD) in all vaccinated animals by Day 63, 7 days after challenge. Placebo controls had moderate to high levels (range 2E+09 – 8.4E+03 gene copies/mL) of SARS-CoV-2 present in nasal swab samples for the duration of the study.

In the lungs (bronchoalveolar lavage, BAL) of SC-Oral-Oral NHP, gRNA also decreased rapidly, with the geometric mean showing a ∼2 log decrease in vaccinated NHP compared to placebo NHP at Day 57, just one day after challenge (Fig. 2C and D, Supplementary Table 4).

In the group receiving an SC and oral boost (SC-SC-Oral), SARS-CoV-2 gRNA in nasal swab samples was also reduced similarly to that seen in SC-Oral-Oral vaccinated primates, with viral gRNA decreasing to levels that were very low or below the LOD in all vaccinated animals by Day 63 (Fig. 2E and F; Supplementary Table 3). In the lungs of SC-SC-Oral NHP, gRNA also showed a ∼2 log decrease on Day 57 (Fig. 2G and H; Supplementary Table 4).

### SC prime, oral boost vaccination immediately inhibited viral replication in nasal passages and lung after SARS-CoV-2 challenge

The presence of replicating virus in nasal swab samples was determined by RT qPCR of subgenomic RNA (sgRNA). By Day 60, 4 days post-challenge, sgRNA was below the LOD for two SC-Oral-Oral primates and, starting on Day 61, below the LOD for all primates that received only oral boosts (Figure 3A and B and Supplementary Table 5).

In the lungs of SC-Oral-Oral NHP, sgRNA also decreased as compared to placebo starting at Day 57, and was below the LOD in all by Day 63 (Fig. 3C and D, Supplementary Table 6).

Evidence of replicating viruses in nasal passages also decreased rapidly in SC-SC-Oral NHP and was below the LOD by Day 59 in two primates and in all primates by Day 63 (Fig. 3E and F; Supplementary Table 5); and in the lungs of this group, sgRNA decreased by ∼2 logs compared to placebo control on Day 57, one day after challenge, with sgRNA being below the LOD at Day 63 in 4 of 5 primates, and just above the LOD in the 5^th^ (Fig. 3 G and H; Supplementary Table 6).

Not only was there a rapid decrease of both viral load and replicating viruses in nasal passages and lung, it is notable that there was no growth of viruses following challenge. This implies the presence of pre-existing humoral and cellular immunity resulting in rapid clearance of the virus upon infection.

### Immediate protection of NHP from SARS-Cov-2 challenge may be due to the presence of T cells responsive to both S and N, and rapid viral clearance to activation of memory B cells

Peripheral blood mononuclear cell (PBMC)-derived T-cell responses to the antigens delivered by the hAd5 S-Fusion + N+ETSD vaccine, spike and nucleocapsid, were determined by ELISpot on Day 0 before prime vaccination, on Day 14 (before boost) and on Day 35, one week after the second Day 28 boost. T cells from SC-Oral-Oral vaccinated primates secreted interferon-gamma (IFN-γ) in response to both S and N peptides on Days 14 and 35 (Fig. 4A). Interleukin-4 (IL-4) secretion was very low (Fig. 4B), indicating the T-cell responses were T helper cell 1 (Th1) dominant, as reflected by the IFN-g/IL-4 ratio (Fig. 4C).

**Fig. 4.**
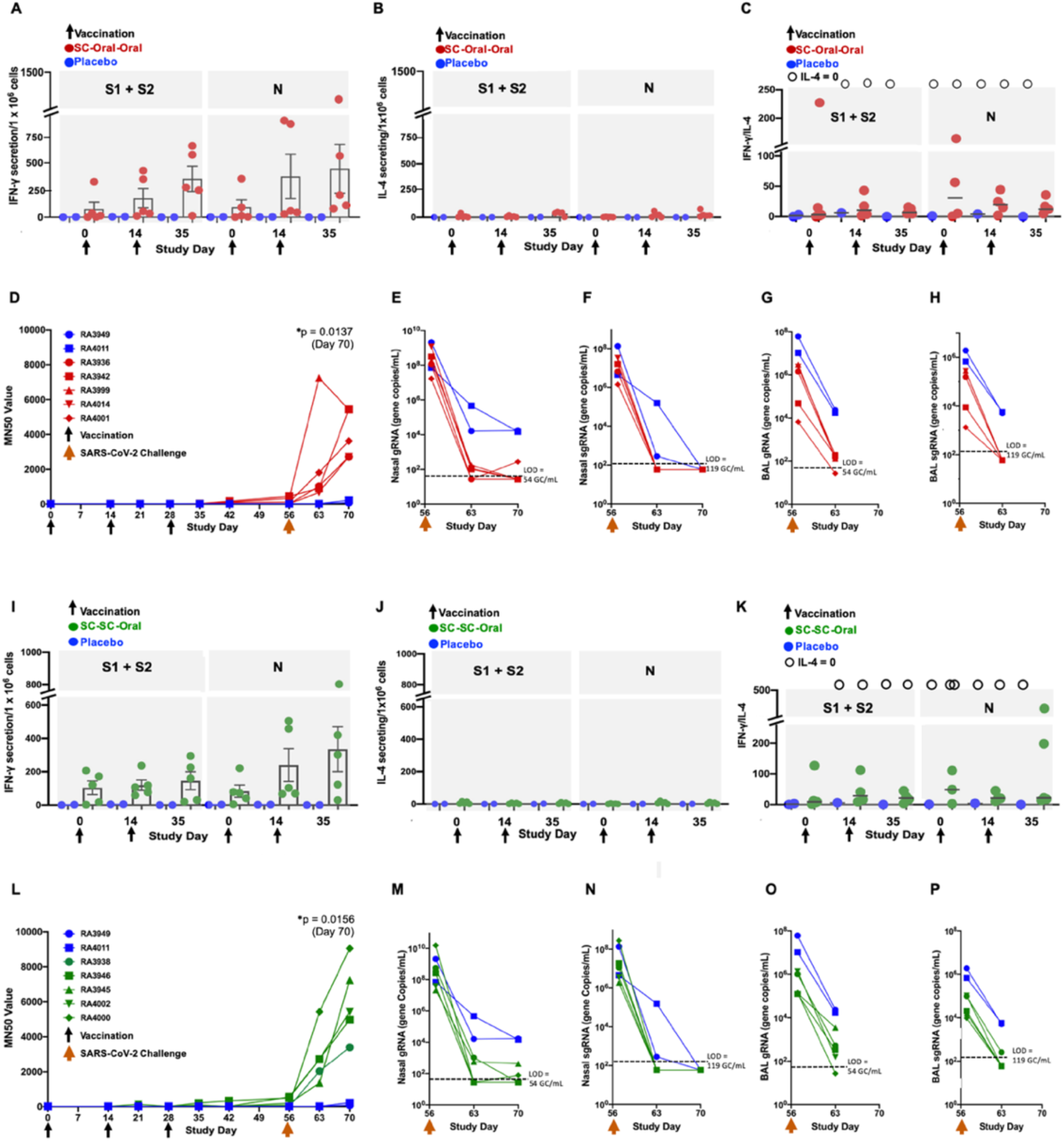
T-cell responses to vaccination and neutralization capability of sera post-SARS-CoV-2 challenge. (A) Interferon-(A) Interferon (IFN-γ) and (B) interleukin-4 (IL-4) secretion by PBMC-derived T cells from SC-Oral-Oral vaccinated NHP in response to spike (S) and nucleocapsid (N) peptides as determined by ELISpot as well as (C) the ratio IFN-γ /IL-4 ratio are shown. Ratios of ‘infinity’ due to undetectable IL-4 are represented as open circles. Cells pulsed with PMA-ionomycin were used as positive controls as described in *Methods* (not presented). Data graphed with mean and SEM. (D) MN50 (dilution factor which SARS-CoV-2 infection of Vero E6 cells is inhibited by 50%) throughout the course of the study is shown; an unpaired, two-tailed Student’s t-test was used to compare MN50s for vaccinated and placebo NHP on Day 70. Nasal gRNA (E) and sgRNA (F) on Days 57, 63 and 70, as well as lung gRNA (G) and sgRNA on Days 57 and 63 are presented. The same data are shown for SC-SC-Oral NHP, including T-cell responses (I-J), MN50 (L), nasal gRNA (M) and sgRNA (N), as well as lung gRNA (O) and sgRNA (P). The level of detection for gRNA was 54 gene copies/mL (GC/mL) and for sgRNA was 119 GC/mL. Half the LOD was used for graphing of data below the LOD.

For sera collected in the post-challenge period, a microneutralization assay (see Methods) was used to assess SARS-CoV-2 neutralization capability as reflected by the ‘MN50’, that is, the serum dilution that correlates to a 50% reduction in viral infectivity as compared to a no-serum control. A rapid increase in neutralization capability of sera for NHP receiving only oral boosts was seen over the two weeks following challenge (Fig. 4D) that mirrored decreases in nasal gRNA and sgRNA (Fig. 4E and F, respectively) and lung gRNA and sgRNA (Fig. 4G and H, respectively). Notably, sera from placebo group primates did not show an increase in neutralizing capability after challenge (Fig. 4D and L), suggesting the existence of memory B cells in the vaccinated group and the absence of such cells in the unvaccinated placebo group. Further studies are needed to confirm this hypothesis.

For SC-SC-Oral vaccinated NHP, findings were very similar for reactive T cells during the pre-challenge vaccination period (Fig. 4I-K) and for neutralization capability in the post-challenge period (Fig. 4L), including the mirroring of decreasing gRNA and sgRNA in nasal passages and lung (Fig. 4M-P).

We hypothesize that the presence of cytotoxic T cells due to vaccination (Fig. 4A-C and I-K) led to the almost immediate decrease in viral replication within the first 24 hours post-challenge (Fig. 4F, H, N, and P), and the continued decreases over the following two weeks that mirrored increases in neutralization capability of sera from vaccinated, but not placebo, NHP reflect the contribution of anti-S-producing memory B cells (Fig. 4D and L).

This study demonstrates that in the rhesus macaque NHP model, subcutaneous prime and oral boost dual-antigen hAd5 S-Fusion + N-ETSD vaccination protects both nasal and lung airways against SARS-CoV-2 challenge. The inhibition of viral replication in nasal passages as evidenced by decreased sgRNA on the first day after viral challenge was notable, as was continuous clearance of virus to levels below detection within 7 days of challenge in all (10/10) animals (Figs. 2 and 3); and, while the rhesus macaque is not a model for the assessment of transmission, these rapid reductions in nasal viral replication are encouraging and support the investigation of the ability of this vaccine to prevent transmission in future studies.

The ability of hAd5 S-Fusion + N-ETSD vaccination to elicit virus-neutralizing anti-S antibodies (Fig. 1) and T cells responsive to both S and N (Fig. 3) - particularly when viewed with the rapid increase in the neutralization capability of sera post-challenge that is likely indicative of the presence of memory B cells - suggests the vaccine establishes broad immunity against severe SARS-CoV-2 infection.

The potential of the hAd5 S-Fusion + N-ETSD SC prime, oral boost vaccine to generate cytotoxic T cells is, in our opinion, a key feature, given the critical role of T cells play in protection from infection in COVID-19 convalescent patients where SARS-CoV-2 specific T cells were identified even in the absence of antibody responses (*27*).

The apparent nearly immediate reduction of viral replication by the hAd5 S-Fusion + N-ETSD vaccination is in contrast to the reported findings for other adenovirus-vectored S-only vaccine NHP studies (*13, 14*), wherein there was evidence of continued viral replication in some animals for at least a day after challenge. Even when challenged with the relatively high titer of 1 ×10^6^ TCID_50_/mL as compared to titers used in some other NHP vaccine studies (*13, 41*), vaccinated animals in our study appeared to be protected from the earliest time point assessed. This rapid protection and clearance was particularly evident in the lung, where both viral load and viral replication were ∼1-2 logs lower than placebo in both vaccinated groups just one day after challenge.

The protection conferred by hAd5 S-Fusion + N-ETSD vaccination of NHPs by SC and oral boost administration particularly reveal the potential for this vaccine to be developed for world-wide distribution, especially in light of the escape variants resistant to antibodies and convalescent plasma, now rapidly spreading throughout the world (*5, 6*). The oral hAd5 S-Fusion + N-ETSD formulation would not require ultra-cold refrigeration like many COVID vaccines currently in development. Dependence on the cold-chain for distribution to geographically remote or under-developed areas causes shipping and storage challenges and will likely reduce the accessibility of the RNA-based COVID-19 vaccines.

Our thermally-stable oral hAd5 S-Fusion + N-ETSD vaccine, due its expression of S and N, also has the potential to act as a ‘universal’ boost to other previously administered vaccines that deliver only S antigens. This use would also be facilitated by cold-chain independence and warrants further exploration.

The hAd5 S-Fusion + N-ETSD vaccine delivered as an SC prime and boost is in Phase 1 clinical trials and the thermally-stable oral vaccine has entered Phase 1 trials as both a prime and boost, and as a boost to an SC prime.

## ACKNOWLEDGEMENTS

We would like to thank the following members of the Battelle group for their contributions in the execution of this study; Chris Cirimotich, Katie Albanese, Phyllis Herr-Calomeni, Bradley Brown, Sara Pfeifer, Carrie Fetzek, Ashley Hay, Andy Puttmann, Amy Allen, and Kevin Coty. We thank Phil Yang of ImmunityBio for his ongoing coordination of project reports for this study. We would also like to thank the DHHS, NIH/NIAID and ASPR/BARDA for funding the study.

## Supplementary Materials

### Materials and Methods

#### The hAd5 S-Fusion + N-ETSD vaccine

To generate the hAd5 S-Fusion + N-ETSD vaccine we utilized the hAd5 [E1-, E2b-, E3-] platform (Fig. S1A) into which a wildtype spike (S) (Fig. S1B) sequence [accession number YP009724390] modified with a proprietary linker peptide sequence was cloned and as well as wildtype nucleocapsid (N) (Fig, S1B) sequence [accession number YP009724397] with a proprietary Enhanced T-cell Stimulation Domain (ETSD) signal sequence to direct translated N to the endosomal/lysosomal pathway. The SARS-CoV-2 S protein is found on the viral surface and its receptor binding domain (RBD) interacts with the host angiotensin-converting enzyme 2 (ACE2) and gains entry to the host cell to initiate infection (*42*). Antibodies against the S RBD are neutralizing, preventing this first step in infection.

The nucleocapsid protein is found in the interior of the virus and is highly conserved and antigenic (*43, 44*). it also plays an important role in T-cell responses (*45, 46*).

The powerful cytomegalovirus (CMV) promoter (*47, 48*) drives expression in the hAd5 construct. The hAd5 S-Fusion + N-ETSD vaccine construct is shown below in Supplementary Fig. S1.

**Fig. S1.**
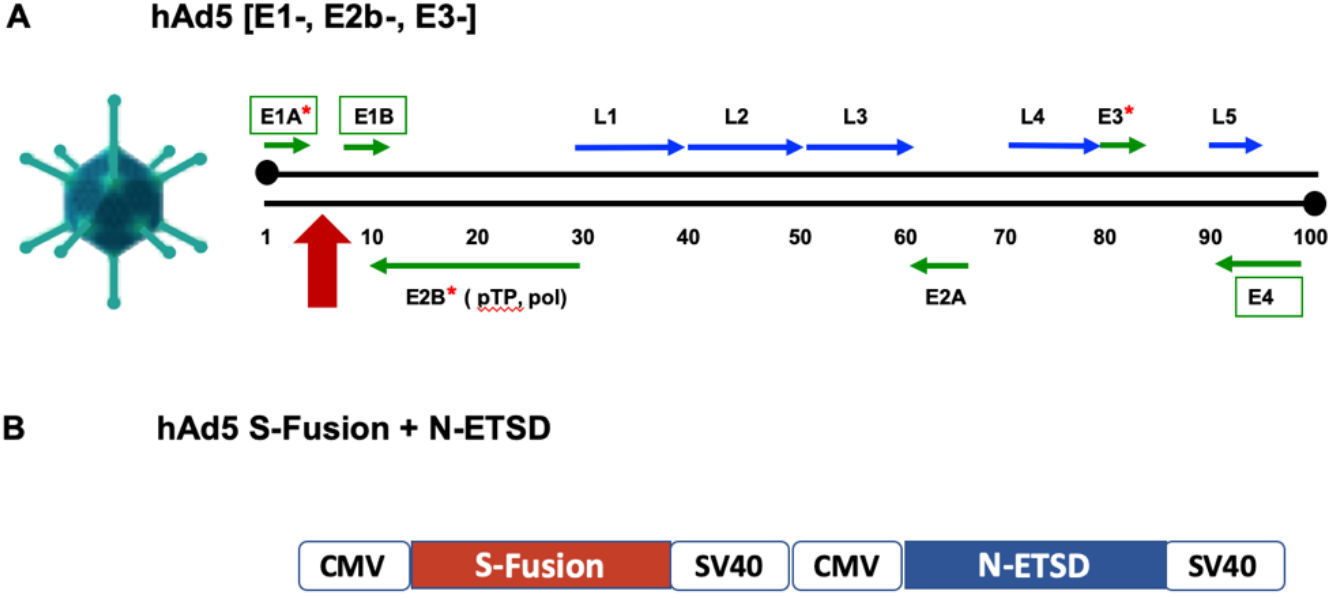
The hAd5 platform and the hAd5 S-Fusion + N-ETSD construct. (A) The human adenovirus serotype 5 vaccine platform with E1, E2b, and E3 regions deleted (*) is shown. The vaccine construct is inserted in the E1 regions (red arrow). (B) The dual-antigen vaccine comprises both S-Fusion and N-ETSD under control of cytomegalovirus (CMV) promoters and with C-terminal SV40 poly-A sequences delivered by the hAd5 [E1-, E2b-, E3-] platform.

#### Transduction of HEK-293T cells and flow cytometry

Relative levels of cell-surface expression of S RBD after transduction of human embryonic kidney (HEK) 293T cells with various S-expressing hAd5 vaccines was determined by flow cytometry. HEK 293T cells (2.5 x 10^5^ cells/well in 24 well plates) were grown in DMEM containing 10% FBS and PSA (100 units/mL penicillin, 100 γ g/mL streptomycin, 0.25 γ g/mL Amphotericin B) at 37°C. Cells were either untransduced or transduced with hAd5 S-WT, S-Fusion, or S-Fusion + N-ETSD viral particles at a multiplicity of infection (MOI) of 10. For detection of spike on Days 1, 2, 3 and 7 after infection, cells were transferred by gently pipetting into medium and labeled with an anti-RBD monoclonal antibody (clone D003 Sino Biological) and F(ab’)2-Goat anti-Human IgG-Fc secondary antibody conjugated with R-phycoerythrin (Thermofisher). Labeled cells were acquired using a Thermo-Fisher Attune NxT flow cytometer and analyzed using Flowjo Software. Results are shown in Supplementary Figure S2.

### The Mouse Study

#### Vaccination of CD-1 mice

CD-1 female mice (Charles River Laboratories) 7 weeks of age were used for immunological studies performed at the vivarium facilities of Omeros Inc. (Seattle, WA). After an initial blood draw, mice were injected with either hAd5 spike wildtype (S-WT) (n = 3) or vaccine candidate hAd5 S-Fusion + N-ETSD (n = 8) on Day 0 at a subcutaneous (SC) dose of 1 ×10^10^ viral particles (VP). Mice received a second SC vaccine dose on Day 21 and on Day 28, blood was collected *via* the submandibular vein from isoflurane-anesthetized mice for isolation of sera and then mice were euthanized for collection of spleen and other tissues.

#### ELISpot assay for assessment of cytokine secretion by T cells

ELISpot assays were used to detect cytokines secreted by splenocytes from inoculated mice. Fresh splenocytes were used on the same day, as were cryopreserved splenocytes containing lymphocytes. The cells (2-4 x 10^5^ cells per well of a 96-well plate) were added to the ELISpot plate containing an immobilized primary containing an immobilized primary antibody to either IFN-γ (BD catalogue # 512525) or IL-4 (BD catalogue # 511819), and were pulsed with 2 μ g/ml of SARS-CoV-2 S or N peptide pools (JPT Peptide Technologies catalogue # PM-WCPV-S-1 and PM-WCPV-NCAP-1 respectively). Negative and positive controls were cells cultured with media alone or Concanavalin A (ConA, 0.46 μg/ml), respectively. After aspiration and washing to remove cells and media, cytokine secretion was detected by biotinylated antibodies to either IFN-γ (BD catalogue # 551818) or IL-4 (BD catalogue # 551804), followed by labeling with a secondary streptavidin/horseradish peroxidase conjugate (BD catalogue # 557630). Spot development was completed using a peroxidase substrate kit (Vector Labs catalogue # SK-4200). The number of spots per well (2-4 x 10^5^ PBMCs) was counted using an ELISpot plate reader. Numbers for graphing were adjusted to spot-forming cells per 10^6^ PBMCs.

#### ELISA for detection of antibodies in the mouse study

For antibody detection in sera from inoculated mice, ELISAs specific for spike and nucleocapsid antibodies, as well as for IgG subtype (IgG1, IgG2a, IgG2b, and IgG3) antibodies were used. A microtiter plate was coated overnight with 100 ng of either purified recombinant SARS-CoV-2 S-FTD (full-length S with fibritin trimerization domain, constructed and purified in-house by ImmunityBio), SARS-CoV-2 S RBD (Sino Biological, Beijing, China; Cat # 401591-V08B1-100) or purified recombinant SARS-CoV-2 nucleocapsid (N) protein (Sino Biological, Beijing, China; Cat # 40588-V08B) in 100 µL of coating buffer (0.05 M Carbonate Buffer, pH 9.6). The wells were washed three times with 250 µL PBS containing 1% Tween 20 (PBST) to remove unbound protein and the plate was blocked for 60 minutes at room temperature with 250 µL PBST. After blocking, the wells were washed with PBST, 100 µL of diluted serum samples were added to wells, and samples incubated for 60 minutes at room temperature. After incubation, the wells were washed with PBST and 100 µL of a 1/5000 dilution of anti-mouse IgG HRP (GE Health Care; Cat # NA9310V), or anti-mouse IgG_1_ HRP (Sigma; Cat # SAB3701171), or anti-mouse IgG_2a_ HRP (Sigma; Cat # SAB3701178), or anti-mouse IgG_2b_ HRP (Sigma; catalog# SAB3701185), or anti-mouse IgG_3_ HRP conjugated antibody (Sigma; Cat # SAB3701192) was added to wells. For positive controls, a 100 µL of a 1/5000 dilution of rabbit anti-N IgG Ab or 100 µL of a 1/25 dilution of mouse anti-S serum (from mice immunized with purified S antigen in adjuvant) were added to appropriate wells. After incubation at room temperature for 1 hour, the wells were washed with PBS-T and incubated with 200 µL o-phenylenediamine-dihydrochloride (OPD substrate (Thermo Scientific Cat # A34006) until appropriate color development. The color reaction was stopped with addition of 50 µL 10% phosphoric acid solution (Fisher Cat # A260-500) in water and the absorbance at 490 nm was determined using a microplate reader (SoftMax® Pro, Molecular Devices).

#### Calculation of relative μg amounts of antibodies

A standard curve of IgG was generated and absorbance values were converted into mass equivalents for both anti-S and anti-N antibodies. Using these values, we were able to calculate that hAd5 S-Fusion + N-ETSD vaccination generated a geometric mean value of 5.8 µg S-specific IgG and 42 µg N-specific IgG per milliliter of serum.

The analysis of antibodies and subtypes, and ELISpot for IFN-γ and IL-4 secretion ratios are shown in Supplementary Figure S3.

### The NHP study

The study, performed at Battelle Biomedical Research Center (Columbus, Ohio), was sponsored by the Biomedical Advanced Research & Development Authority (BARDA), Office of the Assistant Secretary for Preparedness and Response (ASPR), Department of Health and Human Services (DHHS) and the National Institutes of Health/National Institute for Allergy and Infectious Diseases (NIH/NIAID) (Washington, DC). Battelle is a Public Health Service (PHS) Animal Welfare Assurance approved facility. The study protocol was approved by the Institutional Animal Care and Use Committee (ACUC). All aspects of the animal study protocol were designed to minimize stress in the animals.

#### Dosing and sample collection

A total of 12 naïve rhesus macaques weighing >/= 2.5 Kg and being >2.5 years of age were used in the study. All rhesus macaques were tested and confirmed negative within 45 days of receipt for Mycobacterium tuberculosis, simian immunodeficiency virus (SIV), simian T-lymphotropic virus-I (STLV-1), simian retroviruses 1 and 2 (SRV-1 and SRV-2) via PCR, Macacine herpesvirus I (Herpes B virus), and Trypanosoma cruzi (ELISA and PCR).

We compared two SC injections administered in the center of the back just caudal to the scapular region of 1 x 10^11^ vaccine particles (VP) of hAd5 S-Fusion + N-ETSD on Days 0 and 14 followed by an oral capsule 1x 10^10^ infectious units (IU) of hAd5 S-Fusion + N-ETSD delivered via a feeding tube after a minimum of 4 hours of fasting on Day 28 (SC-SC-Oral, Group 1) to one prime SC injection and two oral boost doses with the same dosages and timing (SC-Oral-Oral, Group 2), as shown in Supplementary Figure S4.

Vaccination Group 1 (SC-SC-Oral comprised 3 male and 2 female, Group 2 (SC-Oral-Oral) 2 male and 3 female, and Group 3 (placebo) 1 male and 1 female randomized NHP.

On Day 42, NHPs were transferred to a BSL-3 facility and on Day 56 they were challenged via the intratracheal (0.5 mL) and intranasal (0.25 mL per nares) routes with a total dose of approximately 1 ×10^6^ TCID_50_ SARS-CoV-2 strain USA-WA1/2020. Nasal and oropharyngeal swabs were collected daily from Day 56 (prior to challenge) through Day 63 (7 days post-challenge) and again on Day 70 (14 days post-challenge). In addition, bronchoalveolar lavages (BALs) were performed on Days 57, 59, 61, and 63.

**Fig. S4.**
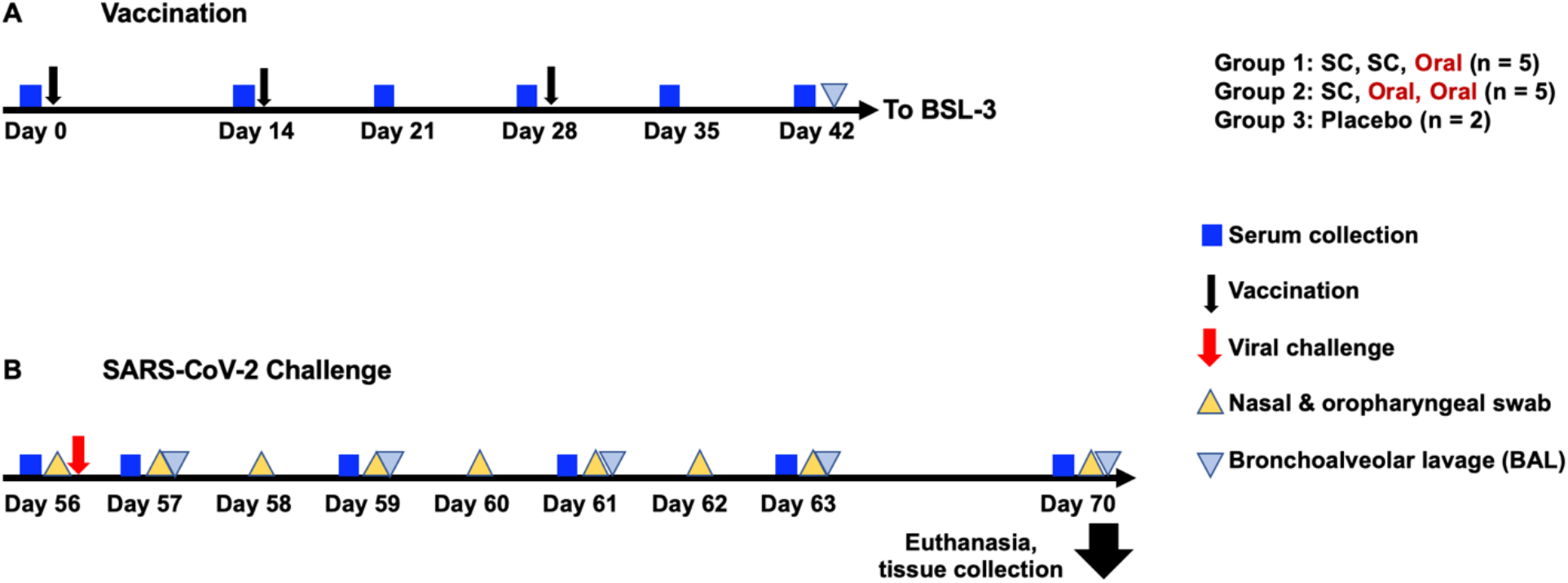
NHP study design. Group 1 NHP (n = 5) were vaccinated by subcutaneous (SC) injection of 1 x 10^11^ VP hAd5 S-Fusion + N-ETSD on Days 0 and 14, and received an oral boost of 1 x 10^10^ IU hAd5 S-Fusion + N-ETSD on Day 28. Group 2 NHP (n = 5) were vaccinated similarly, but both boosts were oral. Sera were collected as indicated (blue box) throughout the study. On Day 42, NHP were transferred to a BSL-3 facility for viral challenge (red arrow) with 1×10^6^ TCID_50_ VP of SARS-CoV-2 intranasally and intratracheally. Nasal and oropharyngeal samples (yellow triangles) and bronchoalveolar lavage (BAL) samples (gray triangles) were collected as indicated. Animals were euthanized on Day 70 and tissues collected for pathological analyses.

#### Clinical signs

NHP in all groups were observed twice daily from study Day -7 until the end of the study on Day 70 for clinical signs, including but not limited to anorexia (weights were taken), hunched posture, lethargy, respiratory distress, activity (recumbent, weak, or unresponsive), convulsions, and other abnormal clinical observations. Blood was collected from a femoral artery or vein, saphenous vein, or appropriate vessel of anesthetized animals at baseline, and Days 14, 21, 28, 35, 42, 56 (challenge Day), 57, 59, 61, 63, and 70 (End Study). Collected blood was used for clinical chemistry and hematological analyses as well as isolation of PBMCs. Body weights are shown in Supplementary Figure S5, hematology in Table S1 and clinical chemistry in Table S2.

#### Statistical analysis

For comparison of animals in groups, One-Way ANOVA was used with an appropriate post-hoc analysis (for example, Tukey’s or Dunnett’s) for specific comparisons. For head-to-head analyses, a two-tailed unpaired Student’s T-test was used. For all statistical analyses, *p <= 0.05, **p < 0.01, ***p<0.001, and ****p<0.0001. All statistical analysis was performed using GraphPad Prism 9 software.

#### ELISA for anti-Spike IgG

IgG against recombinant spike protein in NHP sera or plasma was determined using an Enzyme-Linked ImmunoSorbent Assay (ELISA) wherein 96 well EIA/RIA plates (ThermoFisher, Cat# 07-200-642) were coated with 50 µL/well by a 1 µg/mL solution of purified recombinant SARS-CoV-2-derived Spike protein (S-Fusion; ImmunityBio, Inc.) suspended in coating buffer (0.05 M carbonate-bicarbonate, pH 9.6) and incubated overnight at 4°C. Plates were washed three times with 150 µL of TPBS solution (PBS + 0.05%Tween 20) then 100 µL/well of blocking solution (2% non-fat milk in TPBS) was added and incubated for 1 hour at room temperature (RT). Plasma and serum samples were heat-inactivated at 56°C for 1 hour before use and 1:50 dilutions were prepared in 1% non-fat milk in TPBS. Plates were washed as described above and 50 µL/well of each dilution was added to the plate and incubated at RT for 1 hour. Plates were washed three times with 200 µL of TPBS before addition of 50µL/well of a 1:6000 dilution of HRP-conjugated, cross-absorbed goat anti-monkey IgG (H+L) secondary antibody (ThermoFisher, Cat# PA1-84631) in 1% non-fat milk/TPBS and incubation for 1 hour at RT.

Plates were then washed three times with 200 µL of TPBS and 50 µL of 3,3’,5,5’-tetramethylbenzidine (TMB) substrate (VWR, Cat# 100359-156) was added to each well and incubated at RT for 10 minutes. The reaction was stopped by addition of 50 µL/well of 1N sulfuric acid (H_2_SO_4_). The optical density (OD) at 450 nm was measured using a Synergy 2 plate reader (BioTek Instruments, Inc) and data is analyzed using Prism 8 (GraphPad Software, LLC).

#### cPass™ surrogate assay for determination of the presence of neutralizing antibodies

The presence of neutralizing, anti-spike antibodies in sera from all NHP was determined by assay of sera collected on Days 0, 14, 21, 28, 35 and 42 using the surrogate virus neutralization assay, cPass™ (*49*). The assay is based on assessment of inhibition of binding of the spike receptor binding domain (RBD) to its human host receptor (in the assay, recombinant) angiotensin converting enzyme 2 (ACE2), with inhibition above 20% being correlated with a level of anti-S antibody that is predicted to be neutralizing. All sera samples were diluted 1:30.

#### ELISpot for assessment of cytokine secretion

ELISpot assays were used to detect cytokines secreted by fresh peripheral blood mononuclear cells (PBMCs) isolated from the blood of NHP study animals. PBMCs were isolated from whole blood by standard density gradient centrifugation and frozen in liquid nitrogen until use. PBMCs were thawed and re-suspended in RPMI 10% human AB serum, then pulsed with 2 μg/ml of SARS-CoV-2 S or N peptide pools (JPT Peptide Technologies catalogue # PM-WCPV-S and PM-WCPV-NCAP-1 respectively). Negative and positive controls were cells cultured with media alone or phorbol myristate acetate (PMA, 50 ng/ml) and ionomycin (1 μg/ml), respectively. For IFN-γ assessment, PBMCs were cultured (17 hours at 37°C) in a microtiter plate (Millipore catalogue # MAIPS4510) containing an immobilized primary antibody to capture NHP-specific IFN-γ (MabTech catalogue # 3421M-3-1000). IFN-γ was detected by a secondary antibody to human IFN-γ conjugated to biotin (MabTech catalogue # 3420-6-250). A streptavidin/horseradish peroxidase conjugate (Thermo Fisher catalogue # 21126) was used detect the biotin-conjugated secondary antibody. IFN-γ spot development was completed using a peroxidase substrate kit (Vector Labs catalogue # SK-4200). The number of spots per well (3.5 x 10^5^ PBMCs) was counted using an ELISpot plate reader. IL-4 was assessed using a commercial ELISpot kit (MabTech catalogue # 3410-APW-2) using the manufacturer’s instructions. Numbers for graphing were adjusted to spot-forming cells per 10^6^ PBMCs.

#### Determination of viral load and viral replication post-challenge

RT-qPCR assays were performed to quantify total SARS-CoV-2 RNA copies including genomic RNA using the nucleocapsid protein gene as a target or subgenomic RNA copies that are replication intermediates of the virus using the envelope protein (E) gene as a target. These assays were performed to quantify viral loads following SARS-CoV-2 challenge. RNA was isolated from swabs and bronchioalveolar lavage fluid using the Indispin QIAcube HT Pathogen Kit (Indical Bioscience, Germany) on the QIAcube HT instrument (Qiagen, Germany). The isolated RNA was then evaluated in RT-qPCR using the TaqMan Fast Virus 1-step Master Mix (Thermo Fisher Scientific) on a QuantStudio Flex 6 Real-Time PCR System (Applied Biosystems; Foster City, CA). The primers and probe for total SARS-CoV-2 RNA quantitation were specific to the nucleocapsid protein gene, corresponding to the N1 sequences from the Centers for Disease Control and Prevention (CDC) 2019-Novel Coronavirus (2019-nCoV) Real-Time RT-PCR Diagnostic Panel (https://www.cdc.gov/coronavirus/2019-ncov/lab/rt-pcr-panel-primer-probes.html) except that the probe quencher was modified to Non-Fluorescent Quencher-Minor Groove Binder (NFQ-MGB) (Thermo Fisher Scientific). The primers and probe for the subgenomic RNA quantitation were specific to the E gene subgenomic RNA (Integrated DNA Technologies, Iowa) (*50*). A standard curve comprised of synthetic RNA containing the corresponding target sequence from SARS-CoV-2 isolate WA1 sequence (GenBank Accession Number MN985325.1) (Bio-Synthesis, Inc.; Lewisville, TX) was included on each PCR plate for absolute quantitation of SARS-CoV-2 RNA copies in each sample. Thermocycling conditions were: Stage 1 - 50°C for 5 min for one cycle; Stage 2 - 95°C for 20 sec for one cycle; Stage 3 - 95°C for 3 sec and 60°C for 30 sec for 40 cycles. Data analysis was performed using the QuantStudio 6 software-generated values (total copies per well of each sample) and additional calculations to determine SARS-CoV-2 RNA or subgenomic RNA copies per mL of fluid.

#### Microneutralization assay (MNA)

The neutralizing antibody titer in hAd5 S-Fusion + N-ETSD vaccinated and placebo NHP sera were measured using a Plaque Reduction Neutralization Test (PRNT) carried out in the BSL-4. In brief, the serum samples were heat-inactivated at 56 °C for 90 min, serially diluted two-fold, and pre-incubated with virus stock at 37 °C for 1 hour. The virus/serum mixture was added to 90-100% confluent monolayer Vero E6 cells (BEI, Cat. No. NR-596) in 96-well plates and incubated for 2 days at 37°C with 5% CO_2_. The virus-containing medium was then replaced with 80% acetone for cell fixation. Plates were incubated with an anti-nucleocapsid protein primary antibody cocktail (clones HM1056 and HM1057; EastCoast Bio, North Berwick, ME) for 60 minutes at 37°C. The plates were washed and the secondary antibody (goat anti-mouse IgG Horse Radish Peroxidase (HRP) conjugate; Fitzgerald, North Acton, MA) was added to the wells and the plates were incubated for 60 minutes at 37°C (Battelle Memorial Institute, Patent Number 63/041,551 Pending, 2020). After the plates were washed, the substrate was added and the plates were incubated at 37°C. Stop solution was added and the plates were read for optical density at 405 nm wavelength. Neutralizing activity is defined as at least 50% reduction in signal from the virus only (VC) wells relative to cells control (CC) wells following the formula [(average VC –average CC)/2] + average CC. The median neutralizing titer (MN50) was calculated using Spearman-Kärber analysis method (*51*).

### Supplementary Results

#### Surface expression of spike receptor binding domain (RBDS) in hAd5 S-wildtype (W), hAd5 S-Fusion, or hAd5 S-Fusion + N-ETSD transduced HEK-293T cells

Cell surface expression of S RBD was enhanced with hAd5 S-Fusion compared to hAd5 S-WT, and the dual-antigen hAd5 S-Fusion + N-ETSD construct further enhanced cell surface expression in transduced HEK-293T cells, as shown in Fig. S2.

**Fig. S2.**
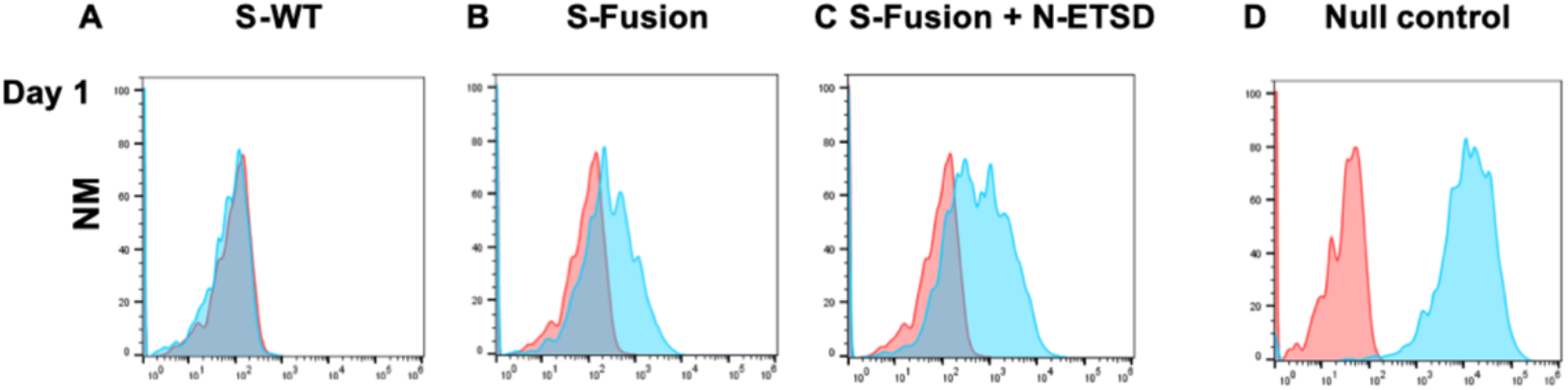
S RBD cell-surface display is enhanced by S-Fusion and S-Fusion + N-ETSD. Cell surface expression of S RBD after transduction of human embryonic kidney (HEK)-293T cells as detected by flow cytometry (normalized to mode, NM) using an anti-RBD antibody with (A) hAd5 S-wildtype (S-WT), (B) hAd5 S-Fusion, (C) hAd5 S-Fusion + N-ETSD, or (D) hAd5 Null control reveals increased surface expression of S RBD by S-Fusion that is further enhanced in S-Fusion + N-ETSD. Untransduced (pink) and transduced (blue).

#### hAd5 S-Fusion + N-ETSD vaccination of CD-1 mice elicits Th1 helper cell dominant antibody and T-cell responses

CD-1 mice vaccinated with the hAd5 S-Fusion + N-ETSD construct produced significantly higher anti-S IgG than mice receiving hAd5 S-WT (Fig. S3A).

**Fig. S3.**
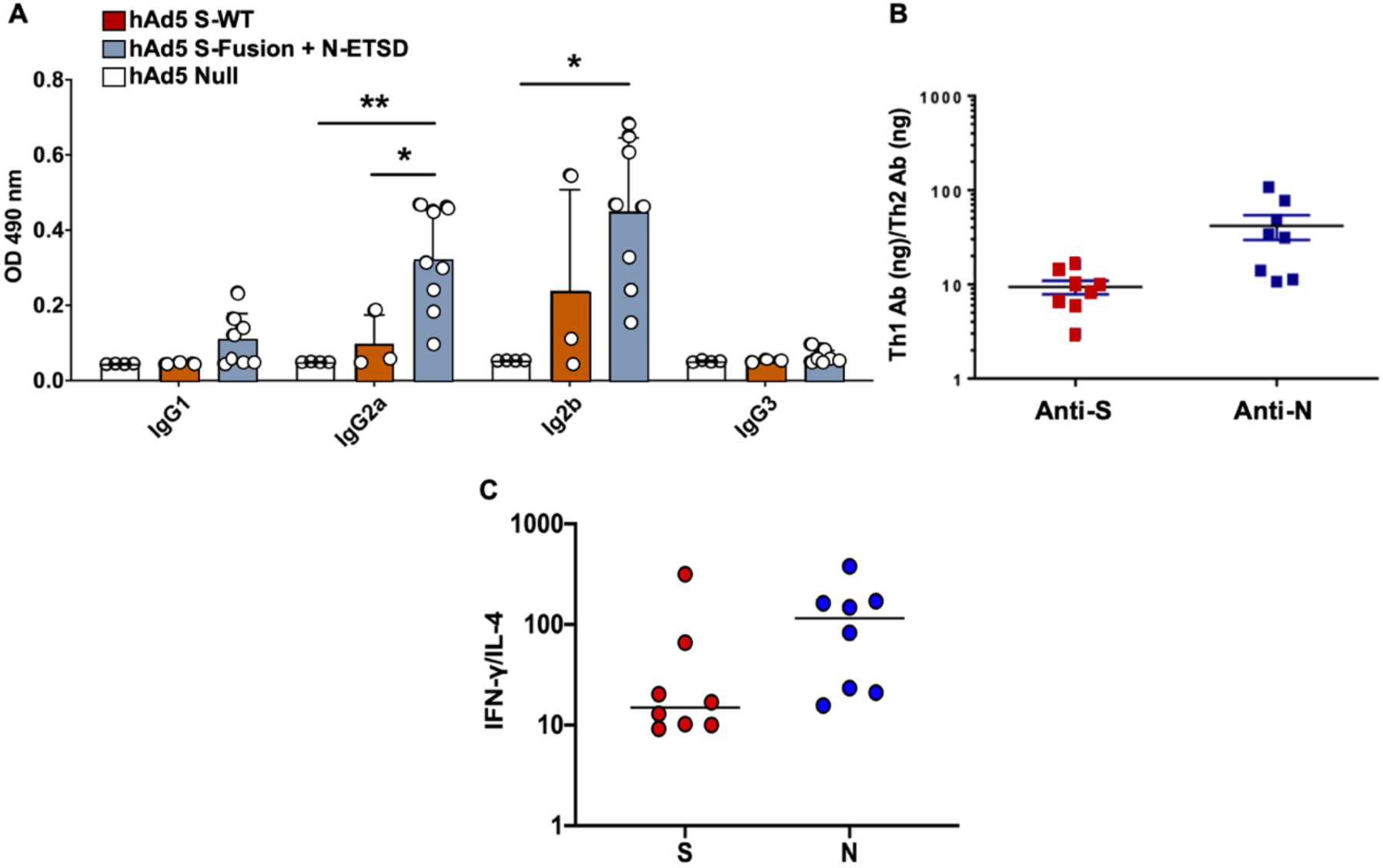
Anti-S IgG and Th1 dominance in hAd5 S-Fusion + N-ETSD vaccinated mice. (A) hAd5 S-Fusion + N-ETSD vaccinated mice produced significantly higher levels of IgG2a and IgG2b anti-S antibodies as compared to mice vaccinated with hAd5 S-WT. Statistical analysis performed using One-way ANOVA and Tukey’s post-hoc comparison of groups. (B) In mice, both anti-S and anti-nucleocapsid (N) antibody responses were Th1 dominant, as were (C) T-cell responses (based on the ELISpot IFN-γ /IL-4 ratio) to both the S and N peptide pools.

### The NHP Study

#### Clinical signs

No clinical signs were noted during the twice daily observations for clinical signs of toxicity due to vaccination and no animals died during the two weeks after one subcutaneous immunization of 1x 10^11^ VP or a week after an oral booster of 1×10^10^ IU of hAd5-S-Fusion+N-ETSD. No gross pathological effect or adverse events were observed and there were no notable changes in body weight (Supplementary Fig. S5).

**Figure S5.**
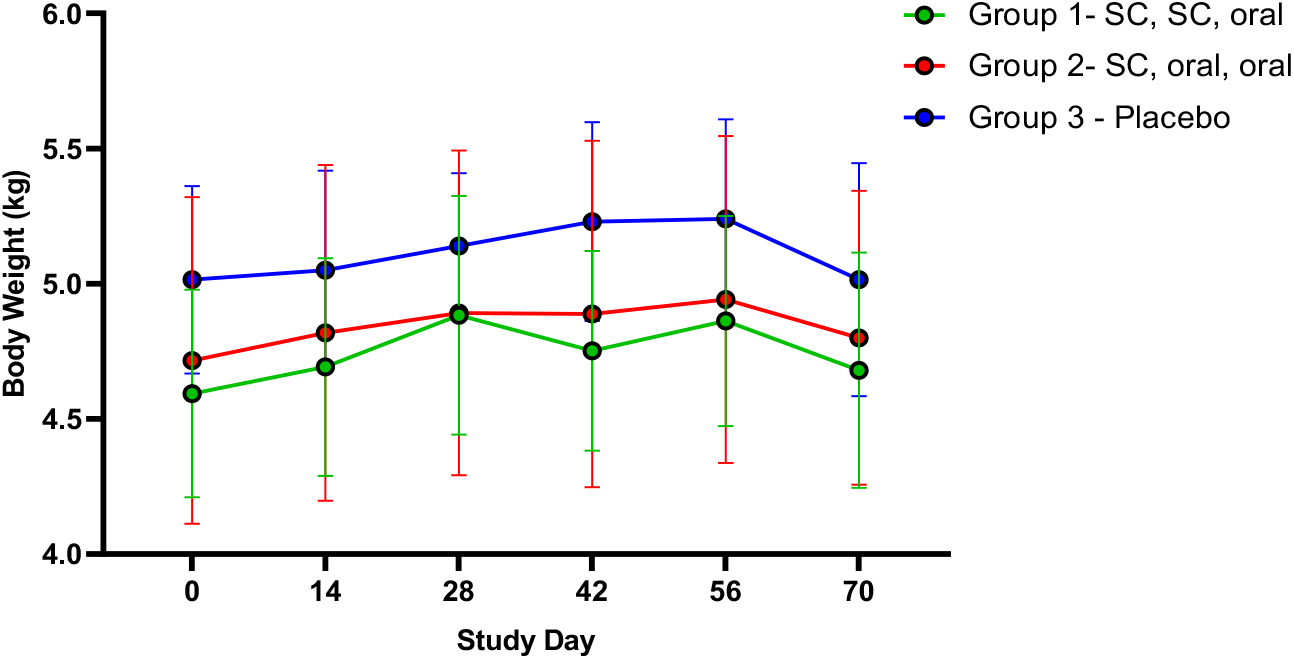
NHP body weights over the course of the study. Body weights are shown in Kg; note y-axis range is 4-6 Kg.

#### Hematology

Hematological parameters evaluated in Group 1 and 2 hAd5 S-Fusion + N-ETSD and Group 3 placebo control animals are presented in Table S1. Only hemoglobin was consistently slightly lower in Group 2 NHP as compared to Group 3 placebo for all post-vaccination time points.

**Table S1.**
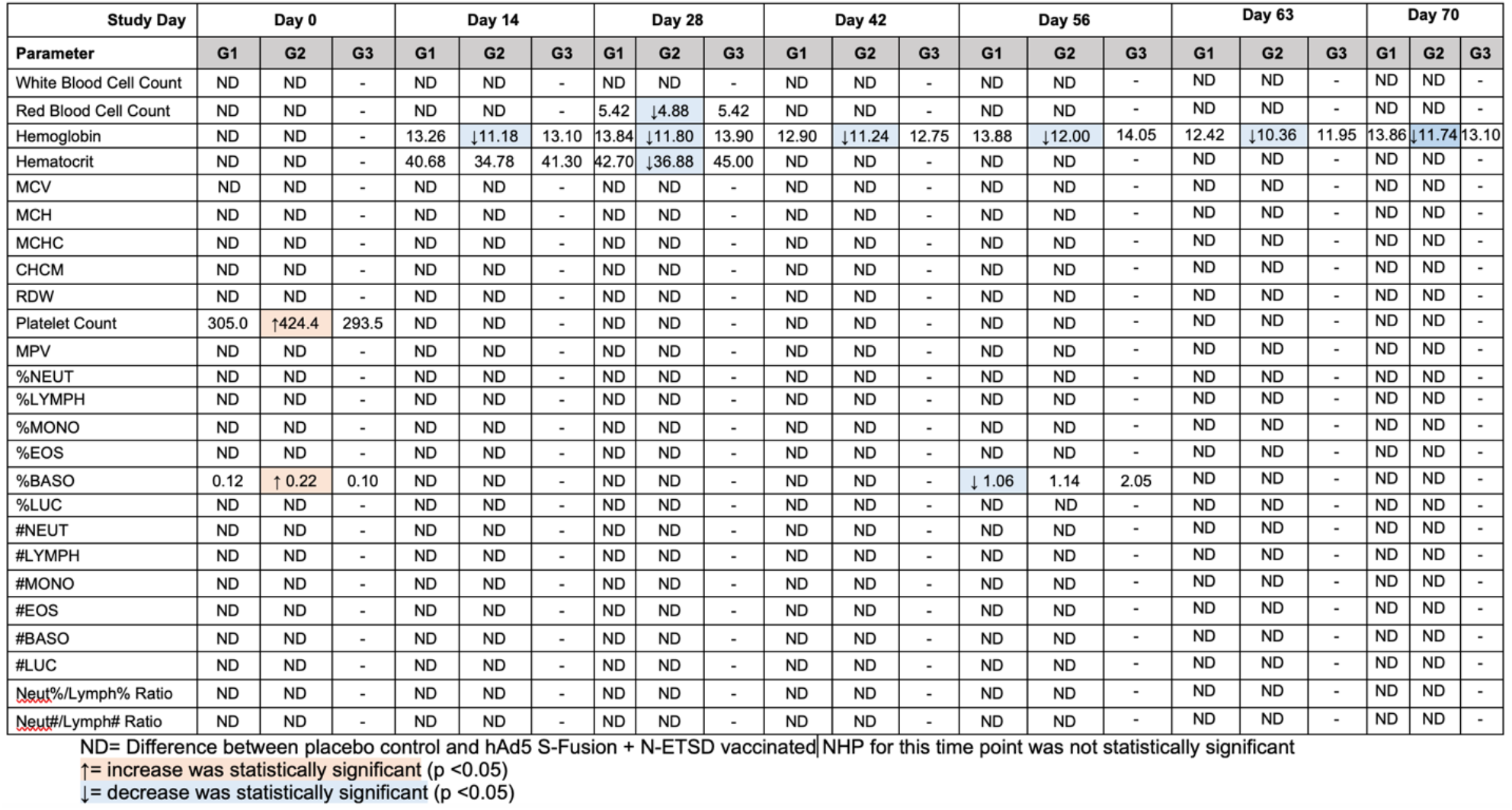
Hematological parameters in hAd5 S-Fusion + N-ETSD vaccinated Group 1 and Group 2 NHP versus placebo Group 3.

#### Clinical Chemistry

Only creatinine was consistently lower in Group 1 NHP on five time points and at two time points for Group 2 (Table S2), but it was lower on Day 0 baseline, indicating this was not vaccine-related. Data in Table S3

**Table S2.**
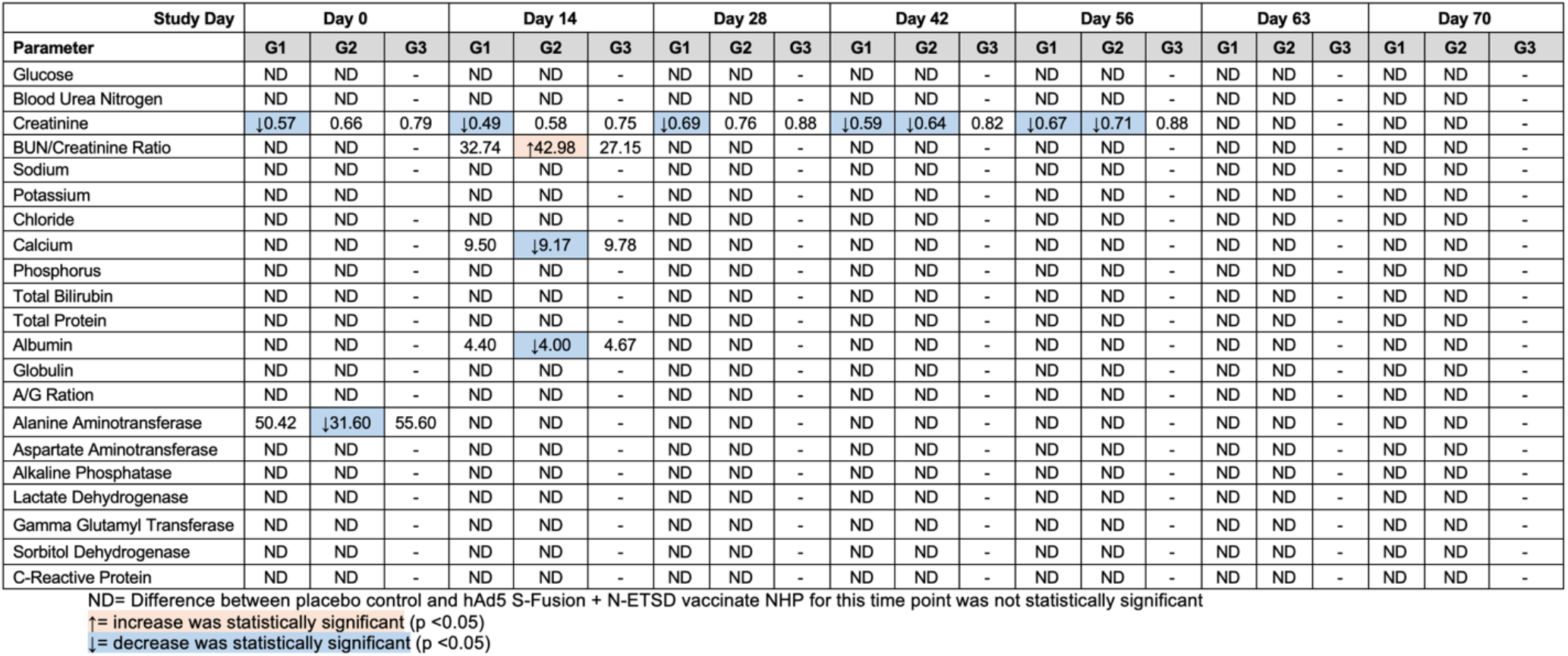
Clinical chemistry for hAd5 S-Fusion + N-ETSD vaccinated Group 1 and Group 2 NHP as compared to placebo Group 3. ND – not significantly different from placebo. Statistical analyses performed using One-way ANOVA.

#### Viral gRNA in nasal swab samples

Viral load was determined in nasal swab samples by RT qPCR of genomic RNA. The data for individual NHP at time point post-challenge are shown in Table S3.

**Table S3.** Gene copies per mL for gRNA as determined by RT qPCR in nasal passages for all groups. Values below the level of detection (LOD) of 54 gene copies per mL (GC/mL) are highlighted in yellow. For calculation of the geometric mean, half the LOD values (27 GC/mL) was used.

| Animal ID | Grp # | Routes | Sex | Day 56 | Day 57 | Day 58 | Day 59 | Day 60 | Day 61 | Day 62 | Day 63 | Day 70 |
| --- | --- | --- | --- | --- | --- | --- | --- | --- | --- | --- | --- | --- |
| RA3938 | 1 | SC-SC-Oral | Male | <LOD | 5.47E+08 | 3.67E+06 | 2.52E+04 | 2.50E+04 | 2.90E+03 | 1.87E+02 | 1.04E+03 | <LOD |
| RA3946 | 1 | SC-SC-Oral | Male | <LOD | 2.59E+08 | 2.04E+07 | 1.38E+06 | 2.46E+05 | 9.01E+04 | 3.29E+03 | <LOD | <LOD |
| RA3945 | 1 | SC-SC-Oral | Male | <LOD | 2.06E+07 | 1.19E+07 | 9.32E+05 | 1.73E+05 | 1.97E+03 | 9.83E+03 | 5.71E+02 | 4.29E+02 |
| RA4002 | 1 | SC-SC-Oral | Female | <LOD | 3.84E+07 | 1.21E+07 | 5.80E+03 | 1.95E+05 | 9.95E+03 | <LOD | <LOD | <LOD |
| RA4000 | 1 | SC-SC-Oral | Female | <LOD | 1.52E+10 | 1.66E+04 | 2.75E+06 | 9.18E+05 | 7.36E+04 | 1.53E+04 | <LOD | 7.79E+01 |
| RA3936 | 2 | SC-Oral-Oral | Male | <LOD | 1.13E+08 | 5.06E+06 | 2.53E+06 | 5.02E+05 | 2.04E+03 | 1.50E+03 | <LOD | <LOD |
| RA3942 | 2 | SC-Oral-Oral | Male | <LOD | 3.13E+08 | 3.59E+06 | 8.15E+05 | 5.62E+03 | 3.32E+02 | <LOD | 1.01E+02 | <LOD |
| RA3999 | 2 | SC-Oral-Oral | Female | <LOD | 1.80E+08 | 2.65E+07 | 1.14E+06 | 3.31E+04 | 2.39E+03 | 1.41E+03 | 1.76E+02 | <LOD |
| RA4014 | 2 | SC-Oral-Oral | Female | <LOD | 1.18E+09 | 4.64E+07 | 2.69E+05 | 3.09E+04 | 1.59E+03 | 1.68E+03 | 1.18E+02 | <LOD |
| RA4001 | 2 | SC-Oral-Oral | Female | <LOD | 1.67E+07 | 4.87E+06 | 3.89E+05 | 3.95E+04 | 5.48E+03 | 1.90E+03 | <LOD | 2.74E+02 |
| RA3949 | 3 | SC-Oral-Oral | Male | <LOD | 2.06E+09 | 4.95E+08 | 4.12E+06 | 7.14E+07 | 7.02E+05 | 2.73E+05 | 1.62E+04 | 1.74E+04 |
| RA4011 | 3 | SC-Oral-Oral | Female | <LOD | 6.95E+07 | 2.38E+07 | 3.00E+07 | 3.44E+05 | 1.56E+05 | 7.19E+05 | 4.61E+05 | 1.47E+04 |

#### Viral gRNA in bronchoalveolar lavage (BAL) samples

Viral load was also determined in BAL samples. Gene copies per mL for individual NHP are shown in Table S4.

**Table S4.** Gene copies per mL for gRNA as determined by RT qPCR in BAL samples for all groups. Values below the level of detection (LOD) of 54 gene copies per mL (GC/mL) are highlighted in yellow. For calculation of the geometric mean, half the LOD values (27 GC/mL) was used.

| Animal ID | Group | Routes | Sex | Day 42 | Day 57 | Day 59 | Day 61 | Day 63 |
| --- | --- | --- | --- | --- | --- | --- | --- | --- |
| RA3938 | 1 | SC-SC-Oral | Male | <LOD | 1.02E+06 | 1.26E+05 | <LOD | 5.24E+02 |
| RA3946 | 1 | SC-SC-Oral | Male | <LOD | 1.25E+05 | 1.04E+05 | 5.43E+04 | 3.31E+02 |
| RA3945 | 1 | SC-SC-Oral | Male | <LOD | 1.33E+05 | 7.50E+04 | 1.22E+05 | 3.54E+03 |
| RA4002 | 1 | SC-SC-Oral | Female | <LOD | 1.40E+06 | 1.59E+04 | 8.74E+03 | 1.55E+02 |
| RA4000 | 1 | SC-SC-Oral | Female | <LOD | 1.44E+05 | 5.00E+03 | <LOD | <LOD |
| RA3936 | 2 | SC-Oral-Oral | Male | <LOD | 1.42E+06 | 3.55E+04 | 8.53E+03 | 1.43E+02 |
| RA3942 | 2 | SC-Oral-Oral | Male | <LOD | 4.80E+04 | 1.40E+02 | <LOD | 1.75E+02 |
| RA3999 | 2 | SC-Oral-Oral | Female | <LOD | 3.00E+06 | 2.28E+06 | 1.06E+06 | 1.31E+02 |
| RA4014 | 2 | SC-Oral-Oral | Female | <LOD | 2.41E+06 | 7.39E+05 | 2.97E+04 | 1.90E+02 |
| RA4001 | 2 | SC-Oral-Oral | Female | <LOD | 6.64E+03 | <LOD | <LOD | <LOD |
| RA3949 | 3 | SC-Oral-Oral | Male | <LOD | 6.11E+07 | 3.44E+06 | 1.16E+05 | 2.35E+04 |
| RA4011 | 3 | SC-Oral-Oral | Female | <LOD | 1.05E+07 | 1.20E+06 | 3.36E+05 | 1.74E+04 |

#### Viral sgRNA in nasal swab samples

Gene copies per mL of subgenomic RNA (sgRNA) for individual NHP are shown in **Table S5**.

**Table S5.** Data for sgRNA in nasal swab samples as determined by RT qPCR passages for all groups. Values below the level of detection (LOD) of 119 gene copies per mL (GC/mL) are highlighted in yellow. For calculation of the geometric mean, half the LOD values (59 GC/mL) was used.

| Animal ID | Group | Routes | Sex | Day 56 | Day 57 | Day 58 | Day 59 | Day 60 | Day 61 | Day 62 | Day 63 | Day 70 |
| --- | --- | --- | --- | --- | --- | --- | --- | --- | --- | --- | --- | --- |
| RA3938 | 1 | SC-SC-Oral | Male | <LOD | 1.12E+07 | 1.27E+05 | <LOD | 3.03E+02 | <LOD | <LOD | <LOD | <LOD |
| RA3946 | 1 | SC-SC-Oral | Male | <LOD | 1.87E+07 | 1.21E+06 | 1.49E+05 | 1.32E+04 | 1.64E+04 | <LOD | <LOD | <LOD |
| RA3945 | 1 | SC-SC-Oral | Male | <LOD | 1.81E+06 | 6.41E+05 | 6.41E+04 | 1.35E+04 | <LOD | 1.07E+03 | <LOD | <LOD |
| RA4002 | 1 | SC-SC-Oral | Female | <LOD | 3.55E+06 | 6.65E+05 | <LOD | 1.45E+04 | <LOD | <LOD | <LOD | <LOD |
| RA4000 | 1 | SC-SC-Oral | Female | <LOD | 2.73E+08 | 4.27E+03 | 2.38E+05 | 6.19E+04 | <LOD | <LOD | <LOD | <LOD |
| RA3936 | 2 | SC-Oral-Oral | Male | <LOD | 6.57E+06 | 4.43E+05 | 1.71E+05 | 2.52E+04 | <LOD | <LOD | <LOD | <LOD |
| RA3942 | 2 | SC-Oral-Oral | Male | <LOD | 1.58E+07 | 3.43E+05 | 1.12E+03 | <LOD | <LOD | <LOD | <LOD | <LOD |
| RA3999 | 2 | SC-Oral-Oral | Female | <LOD | 1.81E+07 | 1.99E+06 | 1.16E+05 | 1.90E+03 | <LOD | <LOD | <LOD | <LOD |
| RA4014 | 2 | SC-Oral-Oral | Female | <LOD | 3.33E+07 | 2.32E+06 | 3.26E+04 | <LOD | <LOD | <LOD | <LOD | <LOD |
| RA4001 | 2 | SC-Oral-Oral | Female | <LOD | 1.42E+06 | 4.97E+05 | 3.84E+04 | 5.98E+02 | <LOD | <LOD | <LOD | <LOD |
| RA3949 | 3 | SC-Oral-Oral | Male | <LOD | 1.33E+08 | 1.84E+07 | 3.21E+05 | 1.49E+06 | 1.23E+04 | 2.73E+03 | 2.86E+02 | <LOD |
| RA4011 | 3 | SC-Oral-Oral | Female | <LOD | 4.47E+06 | 3.81E+06 | 2.48E+06 | 5.88E+04 | 4.40E+04 | 2.04E+05 | 1.56E+05 | <LOD |

#### Viral sgRNA in BAL samples

Gene copies per mL of subgenomic RNA (sgRNA) for individual NHP are shown in Table S5.

**Table S6.** Data for sgRNA in BAL samples as determined by RT qPCR passages for all groups. Values below the level of detection (LOD) of 119 gene copies per mL (GC/mL) are highlighted in yellow. For calculation of the geometric mean, half the LOD values (59 GC/mL) was used.

| Animal ID | Group | Routes | Sex | Day 42 | Day 57 | Day 59 | Day 61 | Day 63 |
| --- | --- | --- | --- | --- | --- | --- | --- | --- |
| RA3938 | 1 | SC-SC-Oral | Male | <LOD | 9.84E+04 | 3.48E+04 | <LOD | 2.58E+02 |
| RA3946 | 1 | SC-SC-Oral | Male | <LOD | 2.06E+04 | 1.12E+04 | 4.57E+03 | <LOD |
| RA3945 | 1 | SC-SC-Oral | Male | <LOD | 1.58E+04 | 1.32E+04 | 1.94E+04 | <LOD |
| RA4002 | 1 | SC-SC-Oral | Female | <LOD | 1.04E+05 | 3.11E+03 | 6.03E+03 | <LOD |
| RA4000 | 1 | SC-SC-Oral | Female | <LOD | 9.63E+03 | <LOD | <LOD | <LOD |
| RA3936 | 2 | SC-Oral-Oral | Male | <LOD | 1.56E+05 | 1.11E+04 | 1.31E+03 | <LOD |
| RA3942 | 2 | SC-Oral-Oral | Male | <LOD | 8.88E+03 | 5.65E+02 | <LOD | <LOD |
| RA3999 | 2 | SC-Oral-Oral | Female | <LOD | 2.81E+05 | 1.38E+05 | 1.47E+05 | <LOD |
| RA4014 | 2 | SC-Oral-Oral | Female | <LOD | 2.74E+05 | 1.19E+05 | 1.24E+04 | <LOD |
| RA4001 | 2 | SC-Oral-Oral | Female | <LOD | 1.32E+03 | <LOD | <LOD | <LOD |
| RA3949 | 3 | SC-Oral-Oral | Male | <LOD | 1.91E+06 | 5.05E+05 | 1.57E+04 | 5.12E+03 |
| RA4011 | 3 | SC-Oral-Oral | Female | <LOD | 6.78E+05 | 9.89E+04 | 8.58E+03 | 5.67E+03 |

## REFERENCES

1. J. T. Wu, K. Leung, G. M. Leung, Nowcasting and forecasting the potential domestic and international spread of the 2019-nCoV outbreak originating in Wuhan, China: a modelling study. The Lancet 395, 689–697 (2020).

2. R. Li et al., Substantial undocumented infection facilitates the rapid dissemination of novel coronavirus (SARS-CoV-2). Science 368, 489–493 (2020).

3. Y. Weisblum et al., Escape from neutralizing antibodies by SARS-CoV-2 spike protein variants. Elife 9, e61312 (2020).

4. Q. Li et al., The Impact of Mutations in SARS-CoV-2 Spike on Viral Infectivity and Antigenicity. Cell 182, 1284-1294.e1289 (2020).

5. S. Cele et al., Escape of SARS-CoV-2 501Y.V2 variants from neutralization by convalescent plasma. medRxiv, 2021.2001.2026.21250224 (2021).

6. H. Tegally et al., Emergence and rapid spread of a new severe acute respiratory syndrome-related coronavirus 2 (SARS-CoV-2) lineage with multiple spike mutations in South Africa. medRxiv, 2020.2012.2021.20248640 (2020).

7. Y. Kou et al., Tissue plasminogen activator (tPA) signal sequence enhances immunogenicity of MVA-based vaccine against tuberculosis. Immunology letters 190, 51–57 (2017).

8. J.-Y. Wang et al., Improved expression of secretory and trimeric proteins in mammalian cells via the introduction of a new trimer motif and a mutant of the tPA signal sequence. Applied Microbiology and Biotechnology 91, 731–740 (2011).

9. P. Sieling et al., Th1 Dominant Nucleocapsid and Spike Antigen-Specific CD4+ and CD8+ Memory T Cell Recall Induced by hAd5 S-Fusion + N-ETSD Infection of Autologous Dendritic Cells from Patients Previously Infected with SARS-CoV-2. medRxiv, 2020.2011.2004.20225417 (2020).

10. K. R. Niazi et al., Activation of human CD4+ T cells by targeting MHC class II epitopes to endosomal compartments using human CD1 tail sequences. Immunology 122, 522–531 (2007).

11. K. Y. Lin et al., Treatment of established tumors with a novel vaccine that enhances major histocompatibility class II presentation of tumor antigen. Cancer research 56, 21–26 (1996).

12. T. C. Wu et al., Engineering an intracellular pathway for major histocompatibility complex class II presentation of antigens. Proceedings of the National Academy of Sciences of the United States of America 92, 11671–11675 (1995).

13. N. B. Mercado et al., Single-shot Ad26 vaccine protects against SARS-CoV-2 in rhesus macaques. Nature 586, 583–588 (2020).

14. N. van Doremalen et al., ChAdOx1 nCoV-19 vaccine prevents SARS-CoV-2 pneumonia in rhesus macaques. Nature 586, 578–582 (2020).

15. A. Amalfitano et al., Production and Characterization of Improved Adenovirus Vectors with the E1, E2b, and E3 Genes Deleted. Journal of virology 72, 926 (1998).

16. M. E. Gatti-Mays et al., A Phase I Trial Using a Multitargeted Recombinant Adenovirus 5 (CEA/MUC1/Brachyury)-Based Immunotherapy Vaccine Regimen in Patients with Advanced Cancer. Oncologist 25, 479 (2019).

17. J. P. Balint et al., Extended evaluation of a phase 1/2 trial on dosing, safety, immunogenicity, and overall survival after immunizations with an advanced-generation Ad5 [E1-, E2b-]-CEA(6D) vaccine in late-stage colorectal cancer. Cancer immunology, immunotherapy: CII 64, 977–987 (2015).

18. E. S. Gabitzsch et al., Novel Adenovirus type 5 vaccine platform induces cellular immunity against HIV-1 Gag, Pol, Nef despite the presence of Ad5 immunity. Vaccine 27, 6394–6398 (2009).

19. M. A. Morse et al., Novel adenoviral vector induces T-cell responses despite anti-adenoviral neutralizing antibodies in colorectal cancer patients. Cancer Immunology, Immunotherapy 62, 1293–1301 (2013).

20. F. R. Jones et al., Prevention of influenza virus shedding and protection from lethal H1N1 challenge using a consensus 2009 H1N1 HA and NA adenovirus vector vaccine. Vaccine 29, 7020–7026 (2011).

21. E. S. Gabitzsch et al., A preliminary and comparative evaluation of a novel Ad5 [E1-, E2b-] recombinant-based vaccine used to induce cell mediated immune responses. Immunology letters 122, 44–51 (2009).

22. E. S. Gabitzsch et al., Induction and comparison of SIV immunity in Ad5 naïve and Ad5 immune non-human primates using an Ad5 [E1-, E2b-] based vaccine. Vaccine 29, 8101–8107 (2011).

23. J. Maruyama et al., Adenoviral vector-based vaccine is fully protective against lethal Lassa fever challenge in Hartley guinea pigs. Vaccine 37, 6824–6831 (2019).

24. S. C. Gilbert, T-cell-inducing vaccines - what’s the future. Immunology 135, 19–26 (2012).

25. N. Le Bert et al., SARS-CoV-2-specific T cell immunity in cases of COVID-19 and SARS, and uninfected controls. Nature 584, 457 (2020).

26. A. Grifoni et al., Targets of T Cell Responses to SARS-CoV-2 Coronavirus in Humans with COVID-19 Disease and Unexposed Individuals. Cell 181, 1489 (2020).

27. T. Sekine et al., Robust T cell immunity in convalescent individuals with asymptomatic or mild COVID-19. Cell 183, 158 (2020).

28. A. J. Greaney et al., Complete Mapping of Mutations to the SARS-CoV-2 Spike Receptor-Binding Domain that Escape Antibody Recognition. Cell Host Microbe, S1931-3128(1920)30624-30627 (2020).

29. Z. Wang et al., mRNA vaccine-elicited antibodies to SARS-CoV-2 and circulating variants. bioRxiv, 2021.2001.2015.426911 (2021).

30. J. Zahradník et al., SARS-CoV-2 RBD in vitro evolution follows contagious mutation spread, yet generates an able infection inhibitor. bioRxiv, 2021.2001.2006.425392 (2021).

31. M. Hellerstein, What are the roles of antibodies versus a durable, high quality T-cell response in protective immunity against SARS-CoV-2? Vaccine: X 6, 100076 (2020).

32. R. Channappanavar, C. Fett, J. Zhao, D. K. Meyerholz, S. Perlman, Virus-specific memory CD8 T cells provide substantial protection from lethal severe acute respiratory syndrome coronavirus infection. Journal of virology 88, 11034–11044 (2014).

33. K. McMahan et al., Correlates of protection against SARS-CoV-2 in rhesus macaques. Nature 10.1038/s41586-020-03041-6, (2020).

34. M. Stewart, S. J. Ward, J. Drew, Use of adenovirus as a model system to illustrate a simple method using standard equipment and inexpensive excipients to remove live virus dependence on the cold-chain. Vaccine 32, 2931–2938 (2014).

35. J. Holmgren, C. Czerkinsky, Mucosal immunity and vaccines. Nature Medicine 11, S45–S53 (2005).

36. F. Xiao et al., Evidence for Gastrointestinal Infection of SARS-CoV-2. Gastroenterology 158, 1831-1833.e1833 (2020).

37. O. Gallo, L. G. Locatello, A. Mazzoni, L. Novelli, F. Annunziato, The central role of the nasal microenvironment in the transmission, modulation, and clinical progression of SARS-CoV-2 infection. Mucosal Immunol, 1–12 (2020).

38. H. Q. Yu et al., Distinct features of SARS-CoV-2-specific IgA response in COVID-19 patients. Eur Respir J 56, 2001526 (2020).

39. L. Moreno-Fierros, I. García-Silva, S. Rosales-Mendoza, Development of SARS-CoV-2 vaccines: should we focus on mucosal immunity? Expert Opinion on Biological Therapy 20, 831–836 (2020).

40. C. W. Tan et al., A SARS-CoV-2 surrogate virus neutralization test based on antibody-mediated blockage of ACE2-spike protein-protein interaction. Nat Biotechnol 38, 1073–1078 (2020).

41. L. Feng et al., An adenovirus-vectored COVID-19 vaccine confers protection from SARS-COV-2 challenge in rhesus macaques. Nature communications 11, 4207 (2020).

42. A. C. Walls et al., Structure, Function, and Antigenicity of the SARS-CoV-2 Spike Glycoprotein. Cell 181, 281-292.e286 (2020).

43. W. Zeng et al., Biochemical characterization of SARS-CoV-2 nucleocapsid protein. Biochemical and biophysical research communications, S0006-0291X(0020)30876-30877 (2020).

44. S. Srinivasan et al., Structural Genomics of SARS-CoV-2 Indicates Evolutionary Conserved Functional Regions of Viral Proteins. Viruses 12, 360 (2020).

45. V. K. Shah, P. Firmal, A. Alam, D. Ganguly, S. Chattopadhyay, Overview of Immune Response During SARS-CoV-2 Infection: Lessons From the Past. Front Immunol 11, 1949 (2020).

46. A. Sattler et al., SARS–CoV-2–specific T cell responses and correlations with COVID-19 patient predisposition. The Journal of Clinical Investigation 130, 6477–6489 (2020).

47. J. Schaack et al., Promoter strength in adenovirus transducing vectors: down-regulation of the adenovirus E1A promoter in 293 cells facilitates vector construction. Virology 291, 101–109 (2001).

48. G. W. Wilkinson, A. Akrigg, Constitutive and enhanced expression from the CMV major IE promoter in a defective adenovirus vector. Nucleic Acids Res 20, 2233–2239 (1992).

49. C. W. Tan et al., A SARS-CoV-2 surrogate virus neutralization test based on antibody-mediated blockage of ACE2-spike protein-protein interaction. Nat Biotechnol 38, 1973 (2020).

50. R. Wölfel et al., Virological assessment of hospitalized patients with COVID-2019. Nature 581, 465–469 (2020).

51. M. A. Hamilton, R. C. Russo, R. V. Thurston, Trimmed Spearman-Karber method for estimating median lethal concentrations in toxicity bioassays. Environmental Science & Technology 11, 714–719 (1977).

